# Single amino acid residues control potential-dependent inactivation of an inner membrane *bc*-cytochrome

**DOI:** 10.1101/2022.08.31.506072

**Authors:** Komal Joshi, Chi Ho Chan, Caleb E. Levar, Daniel R. Bond

**Affiliations:** Department of Biochemistry, Molecular Biology and Biophysics; The BioTechnology Institute; Department of Plant and Microbial Biology, University of Minnesota Twin Cities, USA

**Author notes:** **Corresponding author:** Daniel R. Bond. **Email:**.

## Abstract

During extracellular electron transfer, *Geobacter sulfurreducens* constitutively expresses the *bc-*cytochrome CbcL, yet cells containing only this menaquinone oxidase fail to respire above –0.1 V *vs*. SHE. By identifying mutations within *cbcL* that permit growth at higher potentials, we provide evidence that this cytochrome is regulated by redox potential. Strains expressing CbcL^V205A^, CbcL^V205G^, and CbcL^F525Y^ were capable of growth with high potential electron acceptors including Fe(III) citrate, Mn(IV) oxides, and electrodes poised at +0.1 V *vs*. SHE. Electrochemical characterization of wild type CbcL revealed oxidative inactivation of electron transfer above -0.1 V, while CbcL^V205A^, CbcL^V205G^, and CbcL^F525Y^ remained active. Growth yields of CbcL^V205A^, CbcL^V205G^, and CbcL^F525Y^ were only 50% of WT, consistent with CbcL-dependent electron transfer conserving less energy. These data support the hypothesis that CbcL has evolved to rapidly shut off in response to redox potential to divert electrons to higher yield oxidases that coexist in the *Geobacter* membrane.

**TOC image and caption:** *Tunnel diode behavior:* Electron flux from cells utilizing the menaquinone oxidase CbcL is attenuated by increased redox potential, preventing use of this low-efficiency pathway when driving forces are high enough to conserve energy via other oxidases. Single amino acid substitutions eliminate this switch-off effect and allow function at all potentials. 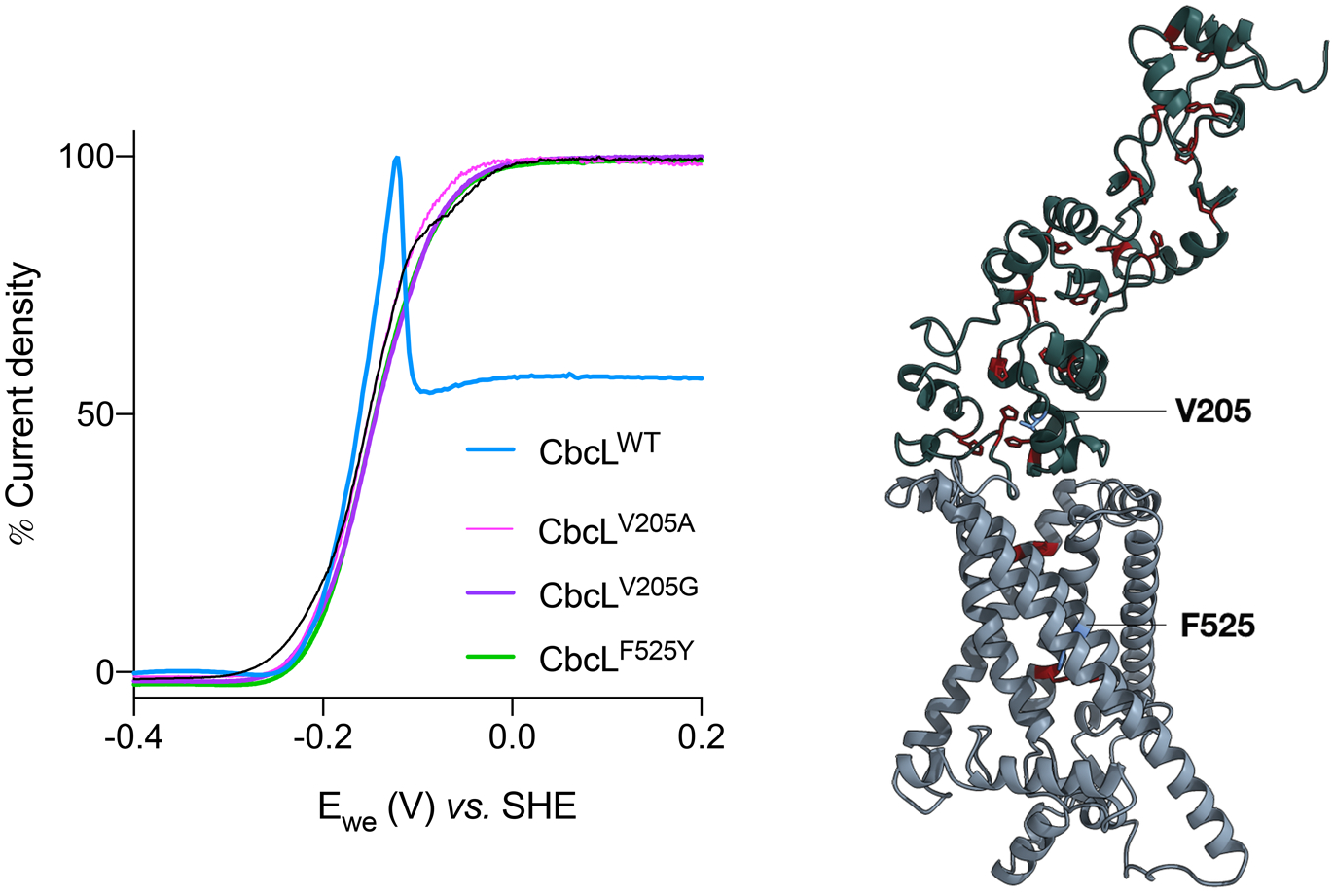

## Introduction

Biological respiration is a universal energy conservation strategy that uses redox reactions to generate transmembrane gradients ^[1,2]^. This mechanism allows cells to vary the number of ions expelled from the cytoplasm in response to the available free energy of electron transfer events. For example, *Escherichia coli* achieves a large H^+^/e^-^ ratio using cytochrome *bo* when oxygen is plentiful, but must switch to cytochrome *bd*–*I* or *bd*–*II* and translocate fewer H^+^/e^-^ as oxygen becomes limiting ^[3]^. Cells can also adjust to declining electron donor availability, such as when *E. coli* switches from the high coupling ratio Hyd-2 to the lower stoichiometry Hyd-1 hydrogenase ^[4]^. Expressing proteins with reduced ion translocation per electron maximizes growth efficiency as available energy decreases.

The dissimilatory metal reducing bacterium *Geobacter sulfurreducens* generates proton motive force while routing electrons from the inner membrane menaquinone pool to cytochromes at the cell’s outer surface, a process termed extracellular electron transfer ^[5,6]^. *G. sulfurreducens* requires different proteins for this task, depending on the redox potential of the extracellular acceptor ^[7–9]^. The seven-heme inner membrane cytochrome *c* ImcH is essential for respiration of the most favorable acceptors, such as Mn(IV) oxides, fresh Fe(III) oxides, and electrodes poised above about -0.1 V *vs*. SHE. Use of ImcH is also correlated with the highest growth rates and yields ^[7,9]^. As redox potential decreases, cells instead require CbcL, a diheme cytochrome *b*/nine-heme *c*–type cytochrome. Below –0.21 V, near the thermodynamic limit of growth with acetate as the donor, the *bc*–type cytochrome complex CbcBA is required for reduction, where cells appear to barely meet maintenance energy requirements. Recent support of this model comes from NMR studies showing the purified *c-*type domain of CbcL has an apparent midpoint potential of -0.194 V, and forms a functional electron transfer complex with the periplasmic cytochrome PpcA (−0.138 V) ^[10]^.

If ImcH, CbcL, and CbcBA are all present when a high potential extracellular acceptor is available, it is thermodynamically possible for electrons to flow through all of them, which would cause use of the less-efficient CbcBA and CbcL pathways. *G. sulfurreducens* solves one part of this problem by only expressing CbcBA when redox potential drops below about -0.15 V. However, all data to date shows that both *imcH* and *cbcL* remain constitutively expressed, regardless of the growth condition ^[8,9,11–13]^. This suggests some kind of post-translational mechanism could exist for preventing use of CbcL at higher potentials.

Cells containing only CbcL (*ΔimcH ΔcbcBA*) do not reduce Fe(III) citrate, electrodes, or Fe(III) oxides when the external redox potential is above -0.1 V. These same cells initiate reduction and growth if potential decreases, then reversibly switch respiration off when potential increases, on a timescale much faster than what can be explained by transcription or translation. Evidence for this effect, where respiration activates and inactivates at a critical potential near -0.1 V is measurable via impedance spectroscopy ^[14]^ and cyclic voltammetry ^[8,9]^, but a mechanism explaining how *Geobacter* controls this change remains unknown.

Devices which decrease in activity as driving force increases are known as tunnel diodes. Succinate dehydrogenases from mitochondria ^[15]^ and aerobic bacteria ^[16]^ resist the thermodynamically favorable fumarate reduction reaction if an increasingly reducing potential is applied, yet catalyze succinate oxidation as potentials are reversed, an effect hypothesized to lend directionality to the TCA cycle. *Escherichia coli* cytochrome *c* nitrite reductase becomes attenuated at reducing potentials, which may prevent partial enzyme turnover before six-electron reduction of the active site occurs ^[17,18]^. Some [NiFe] hydrogenases show strong inactivation by high redox potential, while [FeFe]–hydrogenases can be inactivated at lower potentials ^[19–21]^. Unlike these examples, the tunnel diode-like behavior attributed to *G. sulfurreducens* does not seem to occur in or near the active site of an enzyme. However, all evidence of this gating is based on phenotypes of mutants unable to respire.

In this work, we identify single amino acid substitutions within CbcL that cause gain of function at higher potentials and eliminate tunnel-diode behavior. Cells containing only wild type CbcL could not reduce high potential Mn(IV), Fe(III), and electrode acceptors, but the variants CbcL^V205A^, CbcL^V205G^, and CbcL^F525Y^ reduced all of these compounds nearly as fast as wild type. Voltammetry of cells containing wild type CbcL demonstrated strong attenuation of electron transfer above 0 V vs. SHE, while the single residue variants did not attenuate, and sustained activity above these potentials. When these CbcL variants grew using high potential acceptors, CbcL^V205A^, CbcL^V205G^, and CbcL^F525Y^ achieved lower growth yield compared to wild type cells. This supports a hypothesis where CbcL contributes to a lower H^+^/e^-^ ratio than ImcH, and has evolved an internal redox-dependent switching mechanism to reversibly inactivate and prevent use of its less efficient electron transfer pathway.

## Results

### Enrichment of suppressors in *cbcL* that function at high potential

Strains lacking inner membrane cytochrome ImcH fail to grow in medium containing high potential electron acceptors such as Fe(III) citrate and Mn(IV) oxide ^[7,9]^. Two other inner membrane cytochromes, CbcL and CbcBA, can be used for respiration, but only when electron acceptors are below –0.1 V and –0.21 V, respectively ^[8,9]^. When we incubated Δ*imcH* with Fe(III) citrate, or with poised electrodes (+0.2 V *vs*. SHE), cells did not initially grow, but after a >7 day lag and 3 successive transfers, cultures gained increased rates of reduction. Three Δ*imcH* strains re-isolated from these enrichments reduced Fe(III) citrate significantly faster than the parent. Genome re-sequencing revealed that each contained a single non-synonymous change within GSU0274 (*cbcL*) (Figure 1). In strains isolated from Fe(III) citrate enrichments, valine 205 changed to alanine or glycine, whereas the strain from electrode enrichments replaced phenylalanine 525 with tyrosine.

**Figure 1.**
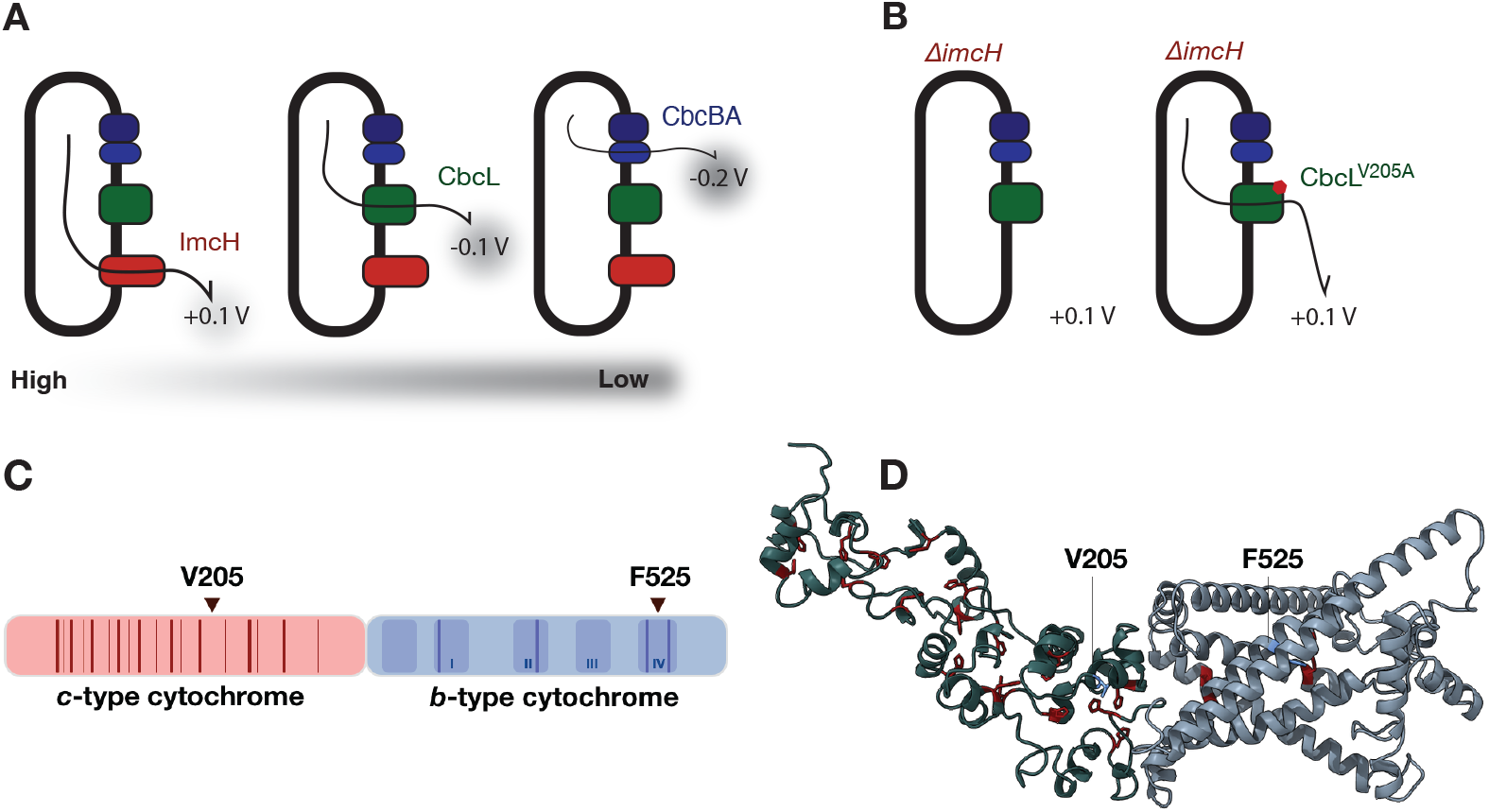
A. Model for electron transfer out of the *Geobacter* cytoplasmic membrane, showing which complex is linked to activity at different redox potentials. B. Strategy for discovery of mutants, using a strain lacking *imcH*, grown under conditions selecting for CbcL function at high potential. C. Cartoon showing the two-domain structure of CbcL, CXXCH motifs indicated by thick red lines, putative coordinating histidines by thin red lines. Transmembrane helices and predicted coordinating histidine residues for the *b-*type domain are in blue. Point mutations discovered within CbcL are indicated on both the primary (C) and AlphaFold2 (D) predicted structure.

### Single amino acid changes in CbcL confer gain of function in Fe(III) citrate

To verify that the *cbcL* point mutations were responsible for the growth phenotype of Δ*imcH* strains, single nucleotides within the genomic copy of *cbcL* were changed in a fresh background where *imcH* was also removed by markerless deletion ^[22]^. These Δ*imcH* CbcL^V205A^, Δ*imcH* CbcL^V205G^, and Δ*imcH* CbcL^F525Y^ strains were then compared to both wild type and the parent Δ*imcH* containing unaltered CbcL^WT^.

The parent Δ*imcH* strain failed to reduce Fe(III), and could not lower the redox potential of the medium. In contrast, the new Δ*imcH* CbcL^V205A^ and CbcL^V205G^ strains reduced Fe(III) citrate at wild type rates to completion (–0.27 V *vs*. SHE) (Figure 2A). The Δ*imcH* CbcL^F525Y^ strain, identified from electrode enrichments, demonstrated slower Fe(III) reduction until redox potential dropped below ∼ 0 V vs. SHE. After this point, Δ*imcH* CbcL^F525Y^ rapidly reduced Fe(III) to the same extent as wild type.

**Figure 2:**
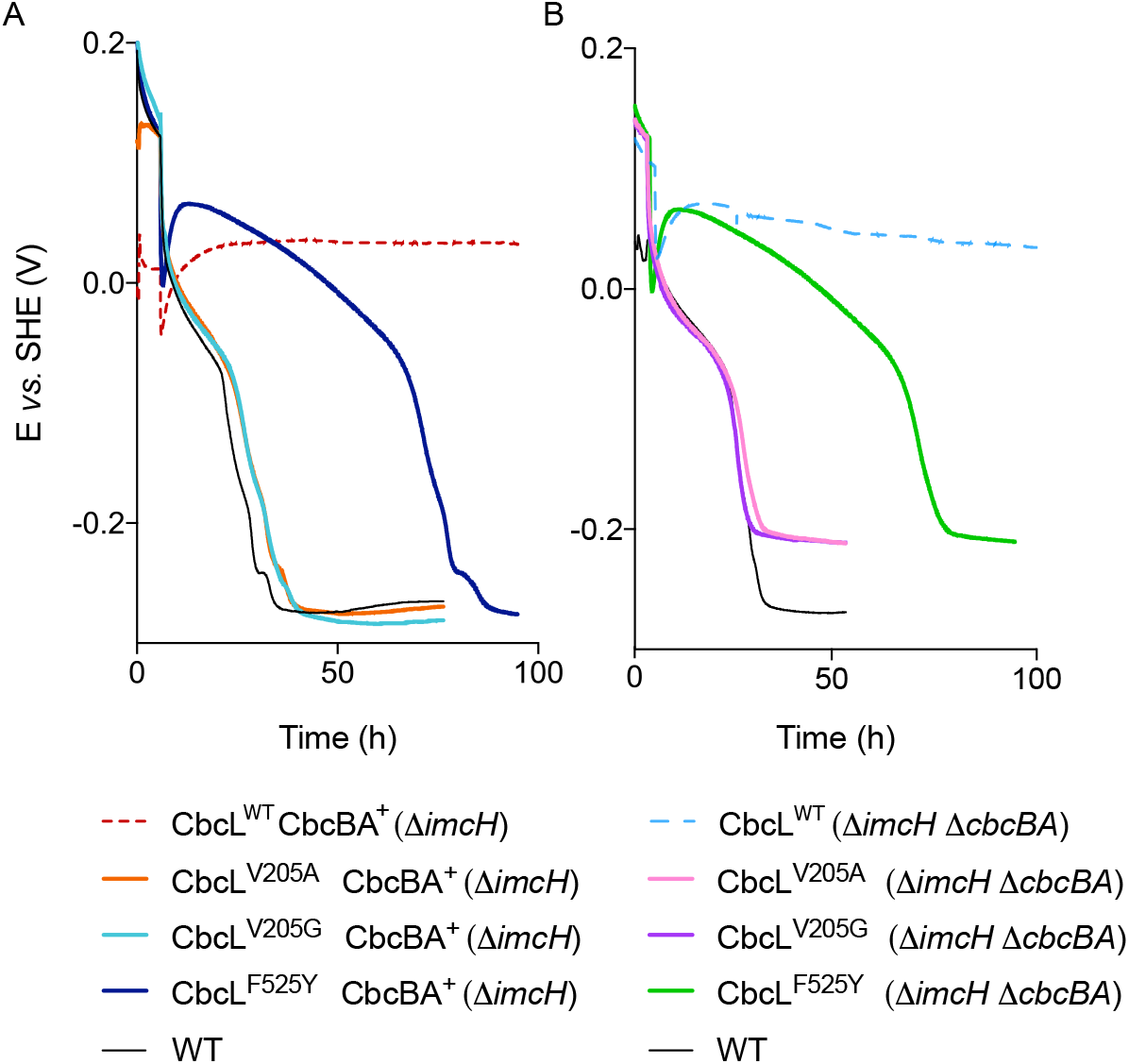
Single amino acid substitutions in CbcL restore growth in high potential Fe(III) citrate. A. Mutants lacking *imcH* with CbcL^V205A^, CbcL^V205G^, and CbcL^F525Y^ can reduce Fe(III) citrate but Δ*imcH* with CbcL^WT^ cannot. B. All strains lacking both *imcH*, and *cbcBA* reduced Fe(III) citrate to the same lower threshold (–0.21 V *vs*. SHE). All experiments were performed in triplicate, and representative data is shown.

A second phenotype linked to CbcL is that it cannot support Fe(III) reduction below -0.21 V vs. SHE, and this is the potential where CbcBA is required to take over electron transfer.

Since mutations in CbcL raised the threshold at which cells could respire, we asked if those mutations altered the lower bound of CbcL activity by removing *cbcBA* ^[9]^. In these double Δ*imcH* Δ*cbcBA* backgrounds, there was no difference between strains containing wild type CbcL and the V205A, V205G, or F525Y CbcL variants. All strains ceased reduction at the same point, or when redox potential reached –0.21 V. Together, this provided evidence that the CbcL residue changes raised the threshold of high potential electron transfer, but left the lower threshold unaltered (Figure 2B).

### CbcL variants also gain the ability to reduce Mn(IV) oxides

While they differ in chemistry and solubility, both Fe(III) citrate and Mn(IV) oxides are high potential acceptors. To test if the CbcL point mutations enabled growth with acceptors other than soluble Fe(III) citrate, strains were inoculated in medium containing 30 mM freshly precipitated Mn(IV) oxide (∼+0.4 V *vs*. SHE) ^[23]^. As previously shown, Δ*imcH* and Δ*imcH ΔcbcBA* mutants failed to reduce any Mn(IV) oxide (Figure 3A, 3B). However, all three *cbcL* variants catalyzed Mn(IV) oxide reduction at wild type levels in both Δ*imcH*, and Δ*imcH ΔcbcBA* backgrounds (Figure 3A, 3B). In addition, initiation of Mn(IV) reduction occurred without any delay in CbcL^F525Y^, unlike Fe(III) citrate where CbcL^F525Y^ demonstrated a significant lag.

**Figure 3:**
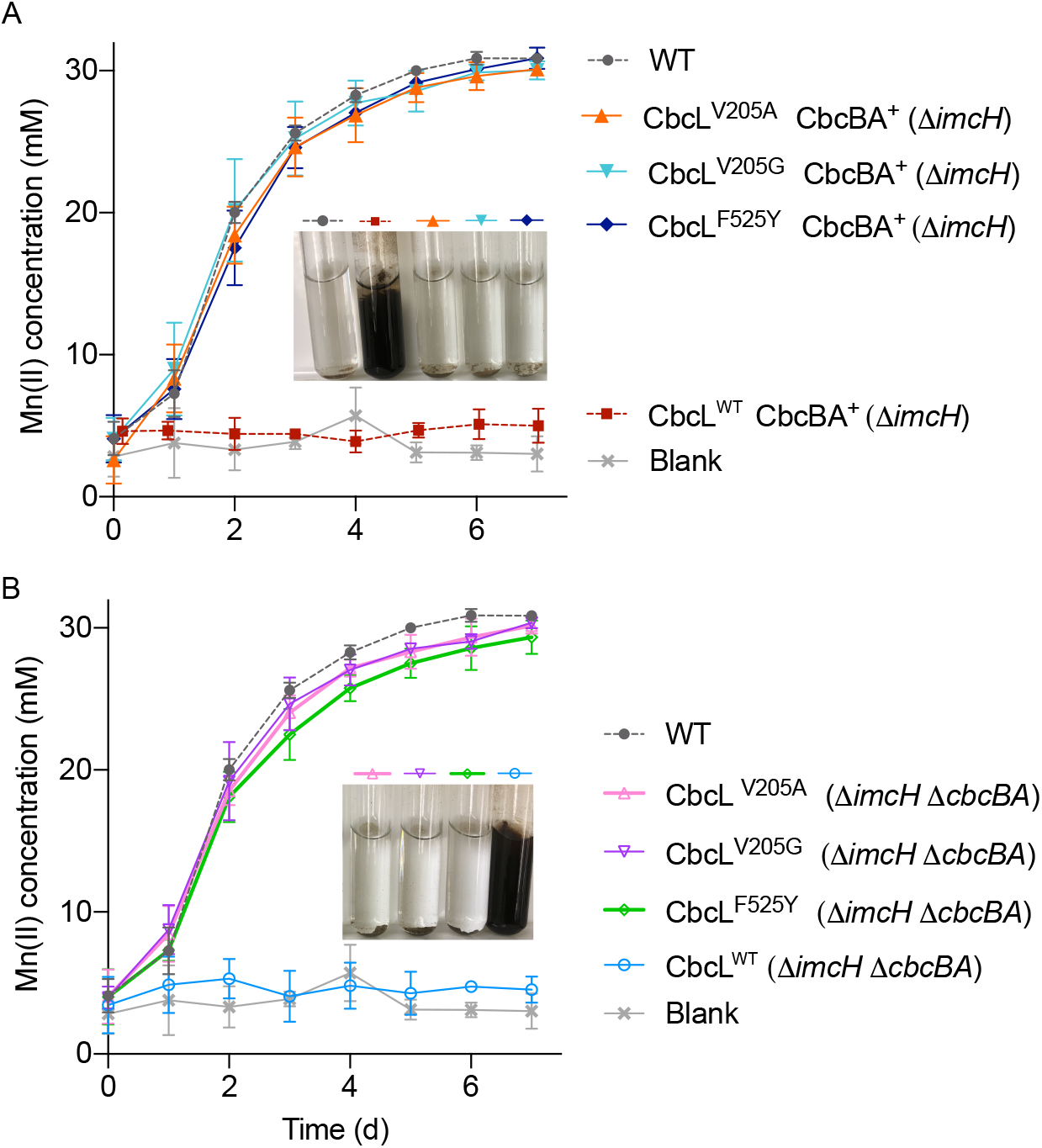
Mutations within *cbcL* enable Mn(IV) oxide reduction. A. Mutants lacking *imcH* with CbcL^V205A^, CbcL^V205G^, and CbcL^F525Y^ can reduce Mn(IV) oxide but Δ*imcH* with CbcL^WT^ cannot. Inset image shows end point of all incubations. B. Double mutants lacking *imcH* and *cbcBA* containing CbcL^V205A^, CbcL^V205G^, and CbcL^F525Y^ reduced all available Mn(IV) oxide to the same extent as WT, but CbcL^WT^ could not. Inset image shows end point of all incubations. Data shows mean and SD of four replicates (n=4).

### CbcL variants also initiate Fe(III) oxide reduction similar to wild type

The redox potential of Fe(III) oxides can be affected by particle size, aging, and pH, but they generally have midpoint potentials lower than Fe(III) citrate and Mn(IV). Strains lacking *imcH* typically show a lag when grown with most Fe(III) oxides, as Fe(II) needs to accumulate in the medium enough to lower potential ^[24]^. When fresh Fe(III) oxides were used, we measured a lag of nearly 5 days before reduction by *ΔimcH* was detected. However, Δ*imcH* strains containing *cbcL* variants did not demonstrate this lag, and instead grew immediately. (Figure 4A, 4C). In Δ*imcH ΔcbcBA* backgrounds, lag was also eliminated in strains containing CbcL^V205A^, CbcL^V205G^, and CbcL^F525Y^, and reduction stopped near the 90% threshold as expected from the lack of *cbcBA*.

**Figure 4:**
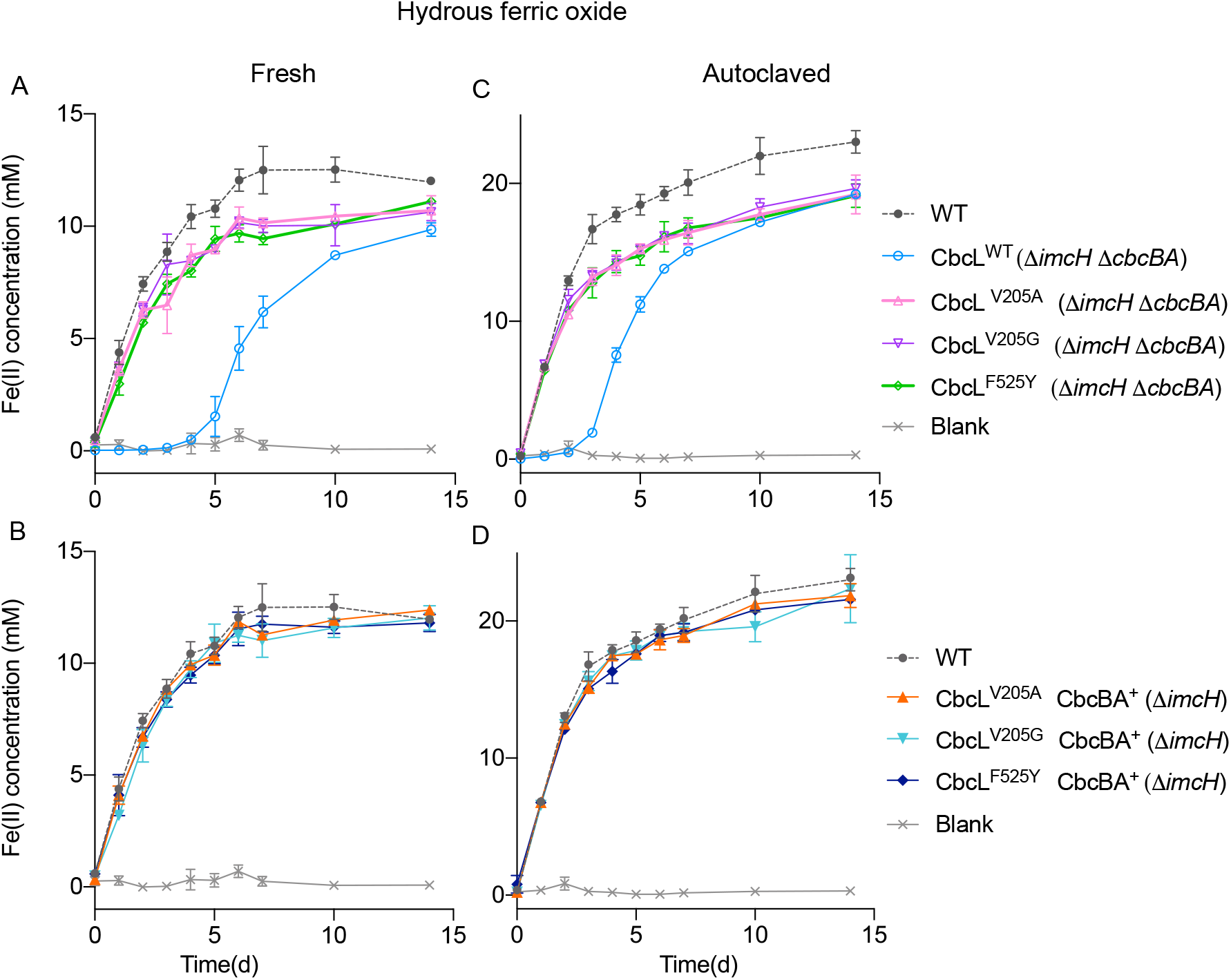
Mutations within *cbcL* eliminate lag with hydrous Fe(III) oxides. Panel A, and B show Fe(III) reduction by single and double mutants containing variants of CbcL, using freshly prepared Fe(III) oxides to create relatively high potential (≥0 V) conditions. Strains containing CbcL^V205A^, CbcL^V205G^, and CbcL^F525Y^ begin reduction immediately, while CbcL^WT^ shows a significant delay. Panel C and D show the effect of autoclaving to lower the potential of Fe(III) oxides. CbcL^WT^ initiated sooner, but CbcL^V205A^, CbcL^V205G^, and CbcL^F525Y^ again had no lag. All strains lacking *cbcBA* only reduced ∼83% of all available Fe(III), showing amino acid substitutions did not alter the lower threshold of reduction by CbcL variants. Experiments were conducted twice with four replicates each. Data is represented as mean ± standard deviation

As an additional test, freshly prepared Fe(III) oxide was autoclaved, an aging treatment known to lower its redox potential and reduce lag of Δ*imcH* strains. As predicted, cells lacking *imcH* initiated Fe(III) reduction sooner with the aged Fe(III), while Δ*imcH* CbcL^V205A^, CbcL^V205G^, or CbcL^F525Y^ still demonstrated no lag (Figure 4A, 4C). Double Δ*imcH ΔcbcBA* CbcL^V205A^, CbcL^V205G^, and CbcL^F525Y^ strains also grew immediately, and stopped at a threshold of ∼83% reduction. These experiments agreed with the model that CbcL variants gained the ability to respire at higher redox potentials, regardless of electron acceptor. The fact that Δ*cbcBA* mutants reduced the same amount of Fe(III), regardless of which CbcL variant was present, also indicated the mutations did not alter the lower bound at which CbcL operated (Figure 4B, 4D).

### Mutations in *cbcL* eliminate oxidative attenuation at high potential electrodes

In addition to high potential metals, mutants lacking ImcH cannot grow on electrodes poised at +0.1 V *vs*. SHE ^[7,24]^. However, Δ*imcH ΔcbcBA* strains containing CbcL^V205A^, CbcL^V205G^, and CbcL^F525Y^ regained the ability to grow at this potential, and achieved similar current densities as wild type within 100 h. Mutants enriched in Fe(III) citrate grew similar to the CbcL^F525Y^ strain which had been enriched on electrodes (Figure 5B).

**Figure 5:**
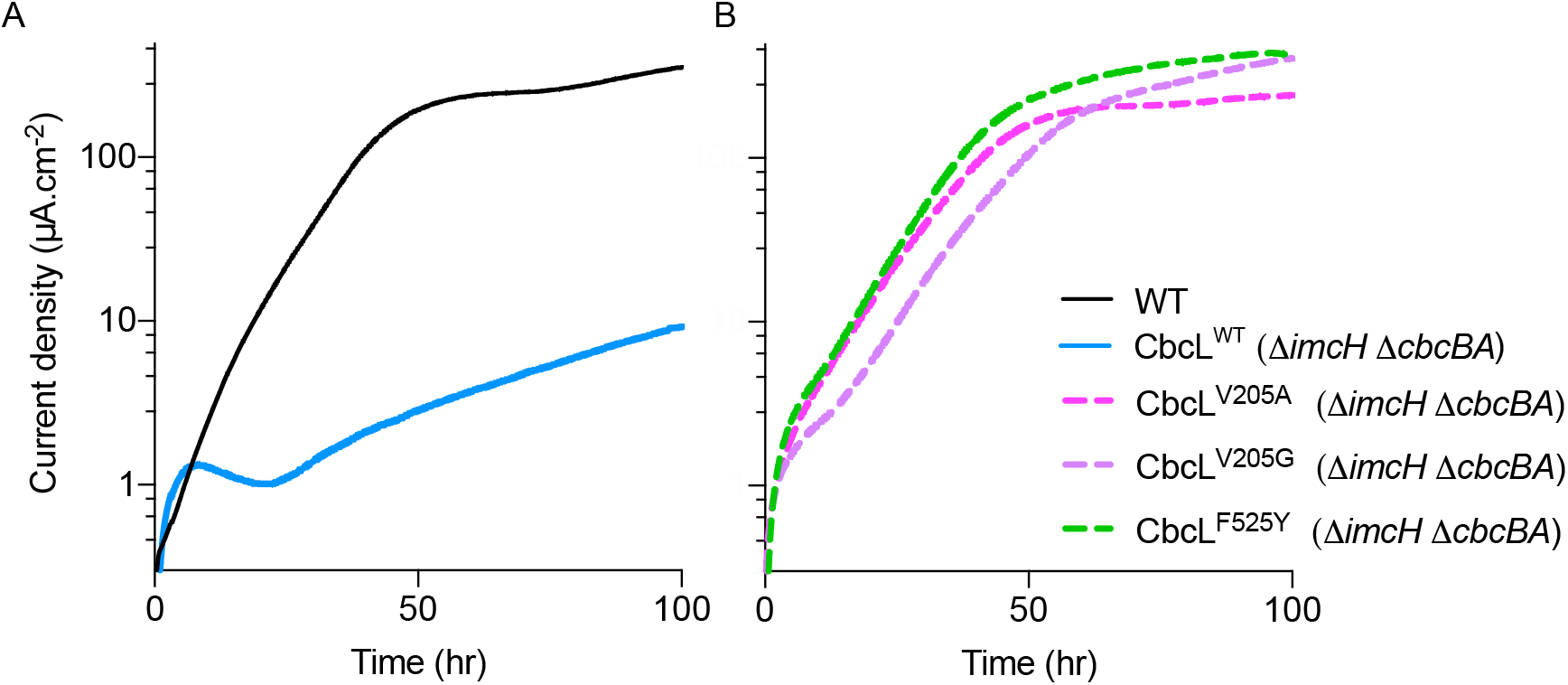
Amino acid substitutions within CbcL restore growth of Δ*imcH ΔcbcBA* cells at +0.1 V *vs*. SHE. A. Current density of WT and cells containing only CbcL^WT^ (Δ*imcH ΔcbcBA*) over time. CbcL^WT^ strain produced less than 10 µA.cm^-2^ after 100 h of incubation while WT reached >400 µA.cm^-2^. B. Variants utilizing CbcL^V205A^, CbcL^V205G^, and CbcL^F525Y^ (Δ*imcH ΔcbcBA*) grew similar to WT at +0.1 V *vs*. SHE. Reactors containing aetate were inoculated with 25% v/v of OD600 ∼0.5 cells grown with fumarate as the electron acceptor. All experiments were performed in triplicate, and representative data is shown.

To investigate changes in voltage-dependent electron transfer, we grew Δ*imcH* Δ*cbcBA* biofilms at –0.1 V vs. SHE, a potential that supports growth of CbcL^WT^, as well as CbcL^V205A^, CbcL^V205G^, and CbcL^F525Y^ (Figure 6A) ^[9]^. In these biofilms containing only CbcL^WT^, current during cyclic voltammetry rose steeply after an onset near –0.2 V, reached a maximum at –0.1 V *vs*. SHE, then dropped sharply even as the redox potential continued to become more favorable (Figure 6A). This “tunnel–diode” effect, where an increase in driving force caused a decrease in current, was reversible, with the midpoint potential of inactivation near –0.11 V (Figure 6B).

**Figure 6:**
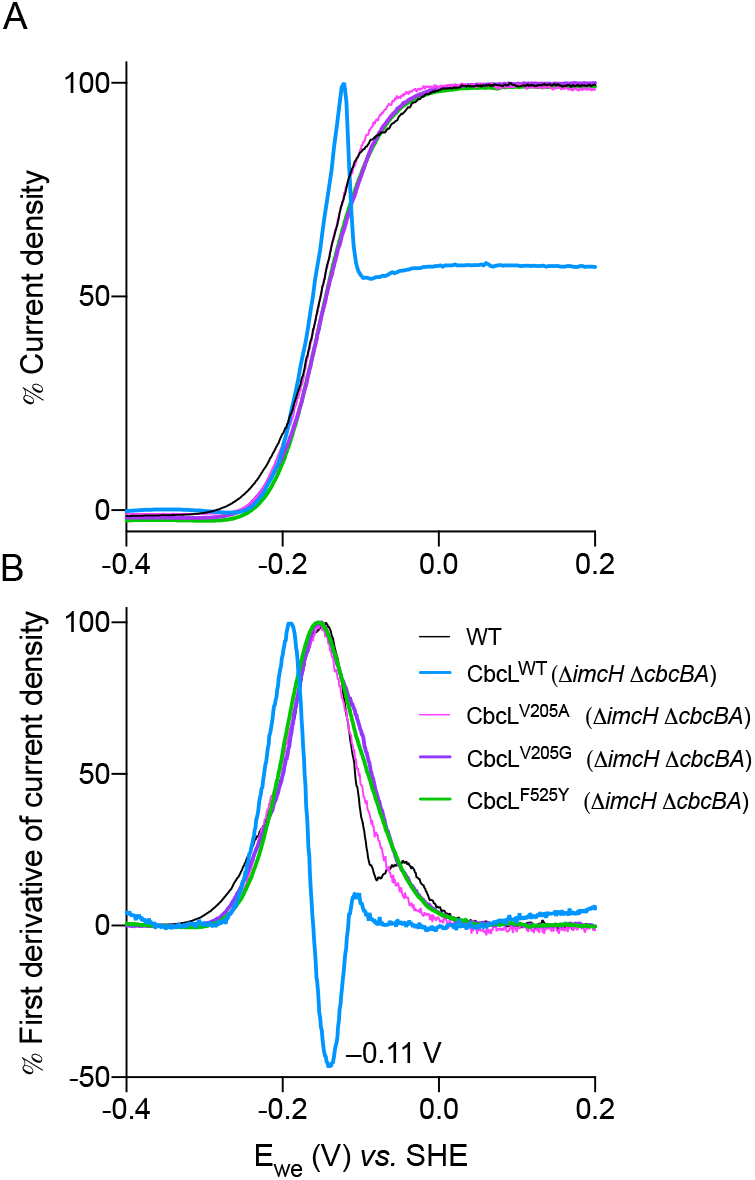
Amino acid substitutions within CbcL eliminate high potential attenuation. A. Cyclic voltammetry (1 mV s^-1^) of *G. sulfurreducens* strains after 96 h of growth. The Δ*imcH ΔcbcBA* biofilms dependent on CbcL^WT^ showed a sharp decrease in current above –0.1 V, while CbcL^V205A^, CbcL^V205G^, and CbcL^F525Y^ did not. The reverse scan for all strains is shown. B. First derivative of scans in A reveal similar midpoints of catalytic features as WT. Data is represented as percentage of limiting current and is representative of triplicate determinations.

In contrast, CbcL^V205A^, CbcL^V205G^, and CbcL^F525Y^ strains showed no such inactivation at potentials as high as +0.2 V vs. SHE. While current produced by CbcL variants continued to rise past the inactivation point of wild type CbcL, the catalytic wave lacked a small feature near -0.05 V vs. SHE that is typical of wild type cells (Figure 6B). This feature is hypothesized to arise from the overlap between CbcL beginning to shut off while ImcH is beginning to function.

While the three variants lost high potential inactivation, they still demonstrated the same low potential characteristics as CbcL^WT^ (Figure 6A).

### CbcL-dependent electron transfer supports lower growth yields

Prior experiments suggested that ImcH, CbcL, and CbcBA support conservation of different amounts of energy during growth with Fe(III) ^[7–9,24]^. However, it has never been possible to directly compare strains using only ImcH to those using only CbcL, as the wild type copy of CbcL is inactivated upon inoculation into medium containing high potential acceptors. When strains containing only ImcH (Δ*cbcL* Δ*cbcBA*) and only CbcL^V205A^, CbcL^V205G^, or CbcL^F525Y^ (Δ*imcH* Δ*cbcBA*) were grown with Fe(III) citrate, and total cells counted using flow cytometry, ImcH-only cells supported the highest cellular yield, or 162 ± 60 % compared to WT (as cells/mM Fe(II), Figure 3.7B). Strains containing only CbcL (CbcL^V205A^, CbcL^V205G^, and CbcL^F525Y^) produced yields of 58.7 ± 28.2 %, 50.1 ± 32.2 %, and 45.5 ± 20.1 % respectively, as compared to WT (Figure 7). While these comparisons are based on mutants within *cbcL*, all three mutants performed similarly, supporting earlier evidence that CbcL and ImcH contribute to different H^+^/e^-^ stoichiometries. It also suggests that in wild type cells, there is a window where both ImcH and CbcL function, which lowers the average yield compared to cells using only ImcH.

**Figure 7:**
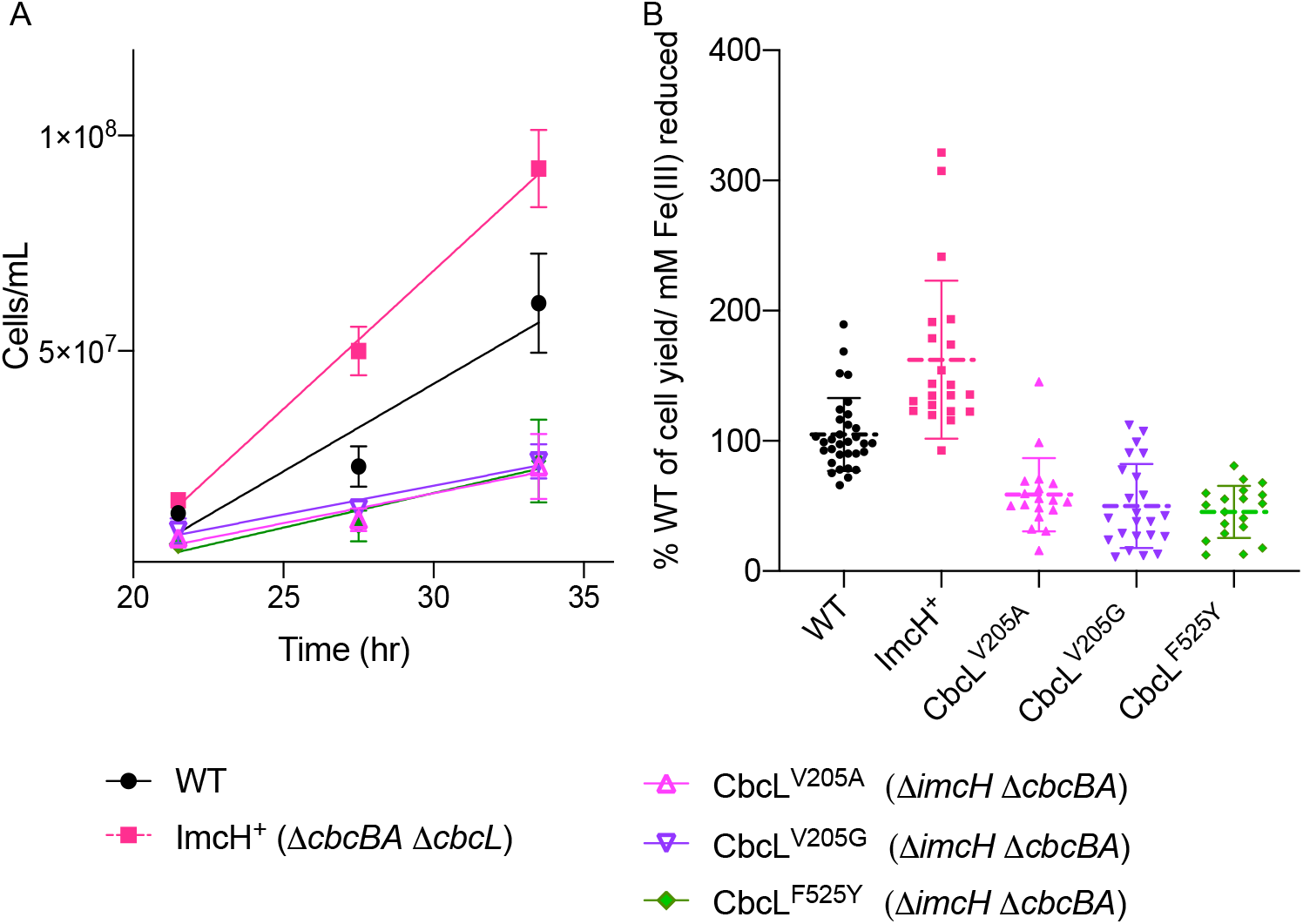
Cell yield of strains utilizing *cbcL* variants is lower than WT. A. Flow cytometry-based cell counts during reduction of Fe(III) citrate. B. Cell yield, calculated as cells per mol Fe(II) produced. Mutants dependent on ImcH (*ΔcbcBA ΔcbcL)* had the highest yield, while strains dependent on CbcL^V205A^, CbcL^V205G^, or CbcL^F525Y^ had lower cell yields than WT. Values ranged from 59 % for CbcL^V205A^, 50 % for CbcL^V205G^, and 46 % for CbcL^F525Y^. Growth experiments were performed three times with eight replicates per run, and data is represented as mean with standard deviation.

## Discussion

While many bacteria utilize different respiratory proteins depending on environmental conditions, most alter their electron transport chain by changing gene expression. Prior work showed that *G. sulfurreducens* constitutively expressed the *bc*-cytochrome *cbcL* ^[12,13]^, yet only appeared to use it for electron transfer when redox potential was low ^[7–9,24]^. The experiments described here show that single mutations within either the membrane or periplasmic domains of CbcL can enable function at high redox potential. This supports the hypothesis that CbcL itself acts as a gate to prevent electron transfer through the protein, to allow use of better options such as ImcH.

Multiple lines of evidence point to redox potential being the primary signal that shuts off CbcL. While cells containing only the wild type copy of *cbcL* were unable to reduce an array of high potential acceptors, all three *cbcL* point mutants gained the ability to grow using multiple forms of Fe(III), Mn(IV), and electrode surfaces (Figures 2, 3, 4, and 5). The data from electrode-grown biofilms, allowing continuous measurement of electron transfer activity across a wide potential window, illustrated this gain of function at higher resolution. Even with mutations in completely different parts of the protein, the cyclic voltammetry response of cells using CbcL^V205A^ and CbcL^V205G^ was nearly identical to CbcL^F525Y^, rising to a stable limiting current with no attenuation as seen in cells using wild type CbcL.

These voltammetry data (Figure 6) also uncovered a possible contradiction, as the lack of growth in Figures 2 and 3 and the electrode growth curves in Figure 5 show electron transfer to be barely detectable above -0.1 V vs. SHE. However, after growth at -0.1 V vs. SHE, raising the potential in a voltammetry sweep triggered only a ∼50% shut-off in electron transfer by the CbcL-dependent biofilm. Why does high potential prevent CbcL function upon inoculation, but in a voltammetry sweep only shut off half of the activity? One possibility is that many of the cell layers in this biofilm were already >5 µm from the surface, where the distance-dependent drop in redox potential created a low-potential region allowing CbcL to continue operating ^[25–29]^. This partial attenuation is also similar to the cathodic tunnel-diode response of nitrite reductase bound to electrodes ^[18]^, suggesting an alternative hypothesis that some fraction of CbcL, held in an ‘on’ state by the consistent driving force supplied by electrodes, becomes insensitive or harder to switch off.

These *cbcL* mutants also allowed testing of a second prediction regarding *Geobacter’s* branched respiratory chain. Prior data using mutants lacking *cbcL* found that cells grown with high potential acceptors had increased growth yields, but the reverse experiment could not be conducted. The ∼50% drop in cell yield measured in all three mutants using CbcL for growth with high potential Fe(III) citrate supports the concept that a lower overall H^+^/e^-^ stoichiometry is achieved when this protein operates.

CbcL is unique in its linkage of a transmembrane *b-*type cytochrome to a soluble *c*-type multiheme cytochrome, without use of any accessory FeS, flavin, NiFe, or molybdopterin subunits. This combination of chemiosmotic modules has not been previously described ^[4]^. The ‘three helix’ *b-*type motif in CbcL, which differs from that in mitochondrial *bc*_1_ and bacterial succinate dehydrogenases ^[30,31]^, allows enzymatic and quinone-based events to occur on opposite sides of the membrane. Fdn formate dehydrogenase, for example, oxidizes periplasmic formate and uses those electrons for cytoplasmic uptake of protons. Similar separations occur in the homologous *b*-type domains of Hyd-1 hydrogenase, Nar nitrate reductase, and Phs thiosulfate reductase ^[32–34]^. In CbcL, this implies reduction of periplasmic cytochromes is coupled to cytoplasmic release of protons, which would not contribute to any redox loops or proton motive force generation. If the quinone-binding site of ImcH is instead at the periplasmic face, as found in similar proteins such as CymA ^[35,36]^, this difference alone could explain the observed differences in growth yields.

How could potential shut off CbcL activity? The highest potential hemes in the purified CbcL *c*-type domain have estimated values of -0.1 V, suggesting full oxidation of the periplasmic region is what deactivates CbcL ^[10]^. Oxidation of hemes could trigger conformational changes that increase heme spacing somewhere within the protein beyond the ∼14 Å boundary required for rapid tunneling, or shift the membrane region to shield the quinone binding site ^[37]^. In AlphaFold2 models^[38]^ (Figure 1), the distance between the membrane *b*-heme and first periplasmic *c*-type heme could be the largest barrier to electron transfer (>10 Å edge-to-edge estimated), and the two domains are connected by a single hinge-like loop. V205 is predicted to be solvent-exposed near this *b-*type and *c-*type domain interface, while F525Y is predicted to introduce a new hydrogen bond between two transmembrane helices. According to a conformational change hypothesis, these mutations stabilize the ‘on’ conformation, but this remains highly speculative until further biochemical and structural work can be conducted.

The CbcL residues found in this work are highly conserved among freshwater *Geobacter* relatives, including the divergent clades containing *G. uraniireducens, G. bemidjiensis*, and *Geomonas spp*.. V205 and F525 also remain invariant across more distant marine groups represented by *Desulfuromonas acetexigens, D. soudanensis*, and *Geoalkalibacter ferrihydridicus*, consistent with these residues being important to CbcL function. Glycine or alanine replaces V205 in strains isolated under unique conditions, such as *Geopsychrobacter electrodophilus* and *Geothermobacter hydrogenophilus*, while dechlorinating *G. lovleyi* and fermentative *Malonomonas* substitute histidine in this position ^[39–42]^. Heterologous expression of these homologs in *G. sulfurreducens*, coupled with voltammetry, could test if these CbcL variants also have altered potential-dependent behavior.

Mechanisms of redox-based regulation can involve a inactivation of electron transfer within a single enzyme, as in [FeFe]– or [NiFe]–hydrogenases ^[20]^, or be mediated by other proteins, such as the thiol-based inactivation of mitochondrial enzymes ^[43]^. Redox changes can even trigger the large-scale reorganization of supercomplexes, as in the photoprotective system of photosystem II ^[44]^. The model emerging for CbcL regulation may represent one of the most fundamental examples that can exist in nature; a wire connecting two pools of reducing equivalents, able to self-inactivate in response to environmental conditions. As respiratory proteins typically evolve out of simpler modular parts, modern enzyme complexes built upon multiheme cytochromes may also possess similar self-regulating abilities, or have the potential to be engineered into switchable bioelectronic devices.

## Materials and Methods

### Bacterial strains and culture conditions

All strains used in this study, including Δ*imcH* suppressor strains, are listed in Table 3.1. Every experiment was initiated by streaking strains from –80 °C frozen culture stocks. All *G. sulfurreducens* strains were cultured in minimal medium (NB) consisting of 0.38 g.L^-1^ KCl, 0.2 g.L^-1^ NH_4_Cl, 0.069 g.L^-1^ NaH_2_PO_4_.H_2_O, 0.04 g.L^-1^ CaCl_2_.2H_2_O, and 0.2 g.L^-1^ MgSO_4_.7H_2_O, 10 mL of trace mineral mix containing 1.5 g.L^-1^ nitrilotriacetic acid (NTA), 0.1 g.L^-1^ MnCl_2_.4H_2_O, 0.5 g.L^-1^ FeSO_4_.7H_2_O, 0.17 g.L^-1^ CoCl_2_.6H_2_O, 0.1 g.L^-1^ ZnCl_2_, 0.03 g.L^-1^ CuSO_4_.5H_2_O, 0.005 g.L^-1^ AlK(SO_4_)_2_.12H_2_O, 0.005g.L^-1^ H_3_BO_3_, 0.09 g.L^-1^ Na_2_MoO_4_, 0.05 g.L^-1^ NiCl_2_, 0.02 g.L^-1^ NaWO_4_.2H_2_O, and 0.10 g.L^-1^ Na_2_SeO_4_, and buffered with 2 g.L^-1^ NaHCO_3_ to a final pH of 6.8 purged with N_2_:CO_2_ (80:20) gas passed over heated copper column to remove trace oxygen.

Nitrilotriacetic acid in trace mineral mix was replaced with HCl to a final concentration of 0.1 M HCl when Fe(III) served as the electron acceptor. For growth on solid medium, 15 g.L^-1^ agar was added to minimal medium containing acetate and fumarate. 20 mM acetate served as the carbon source and electron donor, and 40 mM fumarate as the electron acceptor for routine growth in an anaerobic workstation maintained at 30 °C under N_2_:CO_2_:H_2_ (75:20:5) atmosphere. *Escherichia coli* strains were grown aerobically in lysogeny broth (LB) at 37 °C. 200 µg.mL^-1^ kanamycin was supplemented to *G. sulfurreducens* growth medium, 50 µg.mL^-1^ kanamycin was supplemented to *E. coli* growth medium as necessary. For *G. sulfurreducens*, sucrose to a final concentration of 10 % was added to NB fumarate acetate (NBFA) medium during mutant construction counter-selection step. 55 mM Fe(III) citrate, ∼30 mM amorphous Fe(III) oxide, and ∼30 mM Mn(IV) oxide was used as electron acceptor with 20 mM acetate serving as electron donor for growth with acceptors other than fumarate.

### Mutant construction

Scarless genome modification constructs were designed based on strategy as previously described ^[22]^. Briefly, ∼750 bp flanking *cbcL* (GSU0274) point mutation regions (T614C encoding V205A, T614G encoding V205G, and T1574A encoding F525Y) were amplified using primers listed in Table 3.2. DNA fragments were amplified from genomic DNA extracted from Δ*imcH* suppressor strains 5B2, 5A5, and electrode C respectively. Amplified DNA fragment was digested with XbaI and HindIII, and ligated into the same restriction sites in pk18mobsacB. The ligation product was transformed into UQ950 chemically competent cells. The resulting plasmids were sequence verified before transformation into S17–1 conjugation donor cells. Overnight grown S17-1 donor strain containing the plasmid was conjugated with *G. sulfurreducens* acceptor strain inside an anaerobic chamber on a sterile filter paper placed on an NBFA agar plate. After ∼4 h, cells were scraped from filters, and streaked on NBFA agar plates containing kanamycin. After positive integrants were identified by sequencing, *sacB* counter selection was performed by growing positive integrants on NB fumarate acetate + 10% sucrose plates.

Colonies from sucrose plates were patched on NBFA and NBFA + kanamycin plates to select for antibiotic sensitive, markerless point mutation strains. The mutants were verified by PCR, and Sanger sequencing, and final strains were re-sequenced for off-site mutations via Illumina short read sequencing.

### Enrichment of mutants able to use high potential acceptors

For enrichments on Fe(III) citrate, late exponential phase NBFA grown Δ*imcH* or Δ*imcH::kanR* cultures were inoculated at 1:10 v/v to NB medium containing 55 mM Fe(III) citrate as the sole electron acceptor and 20 mM acetate as the carbon and electron donor. High initial inoculum was used to increase the chances of permissive mutations leading to the enrichment of suppressors. After complete reduction of Fe(III) citrate containing medium, cultures were transferred to fresh Fe(III) citrate containing medium up to five transfers. After the fifth transfer, cultures were streaked for isolation on NB acetate fumarate agar plates. Isolated colonies were picked into 96-well plates with wells containing NBFA medium. Fully grown cultures were transferred by stamping into Fe(III) citrate containing medium. Isolates that cleared Fe(III) citrate medium significantly faster than than of the parent mutant were chosen for further studies.

For electrochemical enrichment, three electrode bioreactors were assembled containing 3 cm^2^ 1500-grit polished graphite working electrode, platinum wire as counter electrode, and saturated calomel electrode as reference electrode. 40 mM acetate served as the electron donor, and 50 mM NaCl was added to maintain osmolarity when fumarate was omitted, and working electrode poised at +0.24 V *vs*. SHE served as the sole electron acceptor. Late exponential phase cells (OD_600_ ≃0.5) grown in NBFA medium were used to inoculate bioreactors with 50 % v/v inoculum size. After current densities of over 50 μA.cm -2 were achieved, bioreactors were taken into an anaerobic chamber, and electrodes were harvested, rinsed once with NBFA medium to remove loosely attached cells, and placed in 10 mL of NBFA medium, then vortexed vigorously to remove attached cells, and were allowed to grow until an OD_600_ ≃0.5 was reached. The electrode enrichment was repeated a total of three times, after which cells from the final electrode were streaked on NBFA agar medium for isolation in anaerobic chamber at 30 °C maintained under N_2_:CO_2_:H_2_ (75:20:5) atmosphere. Five individual colonies were inoculated into NBFA medium. Bioreactors with working electrode poised at +0.24 V *vs*. SHE were inoculated with these cultures after reaching an OD_600_ ≃0.5. Current production was monitored, and strains producing current densities comparable to WT *G. sulfurreducens* were selected for further studies (*7*).

Mutations in selected isolates were identified by sequencing genomic DNA using 50 bp single read or 250 bp paired end reads using the Illumina platform (University of Minnesota Genomics Center). Reads were aligned using breseq (version 0.24rc6) ^[45]^ with *G. sulfurreducens* MN1.

### Fe(III) citrate reduction with constant redox potential measurement

Three electrode bioreactors were assembled with two 1 cm long platinum wire electrodes as working and counter electrodes, and Ag/AgCl electrode (+0.21 V *vs*. SHE) served as a reference electrode. The bioreactors containing 20 mM acetate and 55 mM Fe(III) citrate were continuously flushed with humidified N_2_:CO_2_ (80:20) gas mix to maintain anaerobic conditions. Redox potential was monitored continuously as Fe(III) citrate was reduced using open circuit potential (OCP) method using a 16-channel potentiostat (VMP3, Biologic). Bioreactors were inoculated from NBFA grown cultures when OD_600_ reached ∼0.5 to 1:100 v/v ratio. Before every experiment, platinum electrodes were cleaned electrochemically Platinum was cleaned in 0.5 M H_2_SO_4_ by holding the working electrode at +2.24 V versus SHE, cycling electrode potential between +0.01 and +1.34 V for 20 cycles and stopping at +1.34 V versus SHE. 0.1 mL of samples were taken at regular intervals for Fe(II) concentration measurement using modified ferrozine assay ^[46]^.

### Electrochemical analysis

Three electrode bioreactors were assembled using method as described previously ^[9]^. Briefly, 3 cm^2^ 1500-grit polished polycrystalline graphite served as the working electrode. 2 cm long platinum wire and Ag/AgCl (+0.21 V *vs*. SHE) served as the counter and reference electrode respectively. To maintain anaerobic conditions, reactors were constantly flushed with humidified N_2_:CO_2_ (80:20) gas passed over a heated copper column to remove trace amounts of oxygen. Working electrode was poised at 0 V *vs*. SHE using a VMP3 multichannel potentiostat (Biologic), and a 25 % v/v inoculum of fumarate acetate grown cultures with OD_600_ ≃0.5 was used to initiate the experiments. 40 mM acetate served as the carbon source and electron donor, and poised electrodes served as the sole electron acceptor. Current production was recorded averaged every 120 sec using EC–Lab (Biologic) software.

### Fe(III) oxide reduction

Hydrous ferric oxide, a poorly crystalline form of Fe(III) oxide was synthesized as schwertmannite (Fe_8_O_8_(OH)_6_(SO_4_).nH_2_O) using a rapid precipitation method ^[47]^ by adding 5.5 mL of 30 % hydrogen peroxide to a 10 g.L^-1^ solution of FeSO_4_. Addition of hydrogen peroxide results in precipitation of red-orange schwertmannite mineral. After overnight stabilization of mineral by continuous stirring, schwertmannite solids were washed with DI H 2 O thrice by centrifugation to remove any unreacted FeSO_4_. The washed solids were added to basal medium containing 20 mM acetate supplemented with 0.69 g.L^-1^ NaH_2_PO_4_.H_2_O to prevent crystallization of iron mineral while autoclaving. Schwertmannite mineral is stable in acidic pH but transforms into ferrihydrite mineral with an amorphous XRD–signature at neutral pH ^[47]^. Autoclaving the mineral also changes the properties of the mineral, and has a relatively low redox potential as compared to fresh non-autoclaved mineral ^[48]^. For iron reduction assays, both autoclaved and non-autoclaved forms of hydrous ferric oxide minerals were used. Iron oxide medium was purged with N_2_:CO_2_ (80:20) gas and buffered with NaHCO_3_ to maintain pH at 6.8. Medium containing iron oxide was inoculated with cultures grown in acetate fumarate medium when OD_600_ ≃0.5 at 1:100 v/v. 0.1 mL samples were taken at regular intervals, dissolved in 0.9 Ml 0.5 N HCl, and stored in the dark to prevent photo-oxidation of Fe(II). Fe(II) concentrations were measured using a modified Ferrozine assay.

### Mn(IV) oxide reduction

Mn(IV) oxide was synthesized by mixing 20 mM KMnO_4_ and 3.2 g.L^-1^ NaOH. To it, a 1000 mL solution of 30 mM MnCl_2_.4H_2_O was added ^[49]^. The mix was continuously stirred at room temperature for at least two hours, and then moved to 4 °C with continuous stirring overnight. The continuous stirring, and long incubation helped with mineral formation. The MnCl_2_ and NaOH served to stabilize the pH. The resulting amorphous mineral was washed with DI H_2_O thrice by centrifugation to remove any unreacted KMnO_4_. The Mn(IV) oxide yield was measured by drying out the mineral, and measuring weight. The yield of Mn(IV) oxide mineral used in the experiments was calculated to be ∼0.5 M. The MnO_2_ was diluted to a final concentration of ∼0.3 Mn(IV) oxide in basal medium lacking carbon source. This diluted Mn(IV) oxide containing medium was purged with argon gas passed over a heated copper column to make Mn(IV) oxide medium anaerobic. To bring the final concentration of Mn(IV) oxide to ∼30 mM, 1 mL of Mn(IV) oxide stock solution was added to autoclaved 9 mL *Geobacter* basal growth medium lacking electron acceptor supplemented with 10 mM acetate as carbon source and electron donor. 1 mL of Mn(IV) oxide was added to the tubes immediately before inoculation. Fumarate acetate grown cultures at OD_600_ ≃0.5 were inoculated at 1:100 v/v, and 0.1 mL samples were taken at regular intervals and dissolved in 0.9 mL of freshly prepared 2 N HCl with 4 mM FeSO_4_ for Mn(IV) reduction measurements. The samples were stored in dark before measurement via a modified Ferrozine assay. As the reduction of Mn(IV) by Fe(II) is thermodynamically favorable, the measured Fe(II) concentration can be used to calculate the Mn(II) present in a given sample. Mn(II) concentrations were measured indirectly by measuring Fe(II) concentration by ferrozine assay.

### Flow cytometry analysis and yield calculation

Double filtered medium containing 55 mM Fe(III) citrate and 20 mM acetate was used to count total cells using flow cytometry. Filtered acetate fumarate medium grown cultures (OD_600_ ≃0.5) at 1:100 v/v were used to inoculate Fe(III) citrate containing medium. All the glass wares used in this experiment were rinsed thrice with 0.45 µm filtered DI H_2_O. Samples were taken at regular intervals for cell counting, and for Fe(III) reduction measurement via ferrozine assay.

Cell counting measurements were conducted using a FACS Calibur flow cytometer (BD Biosciences) equipped with 488 nm and 640 nm lasers through a 530/30 filter. Photodiode setting on E00 with Amp Gain of 1 was applied to forward scatter detector, photomultiplier tube (PMT) setting at 415 was applied to collect side scatter data. A threshold value of 90 was set for all measurements. All dilutions were made in filtered NB non-growth medium lacking any carbon sources. Sample flow rate used was 35 ± 5 µL.min^-1^ (medium flow), and samples were taken in 51 s passes. All events were recorded. Population counting was performed using FlowJo software (BD Biosciences). Fe(III) reduction was measured using ferrozine assay, and yield was calculated as total increase in cell counts per mM Fe(III) reduced.

**Table 1:**
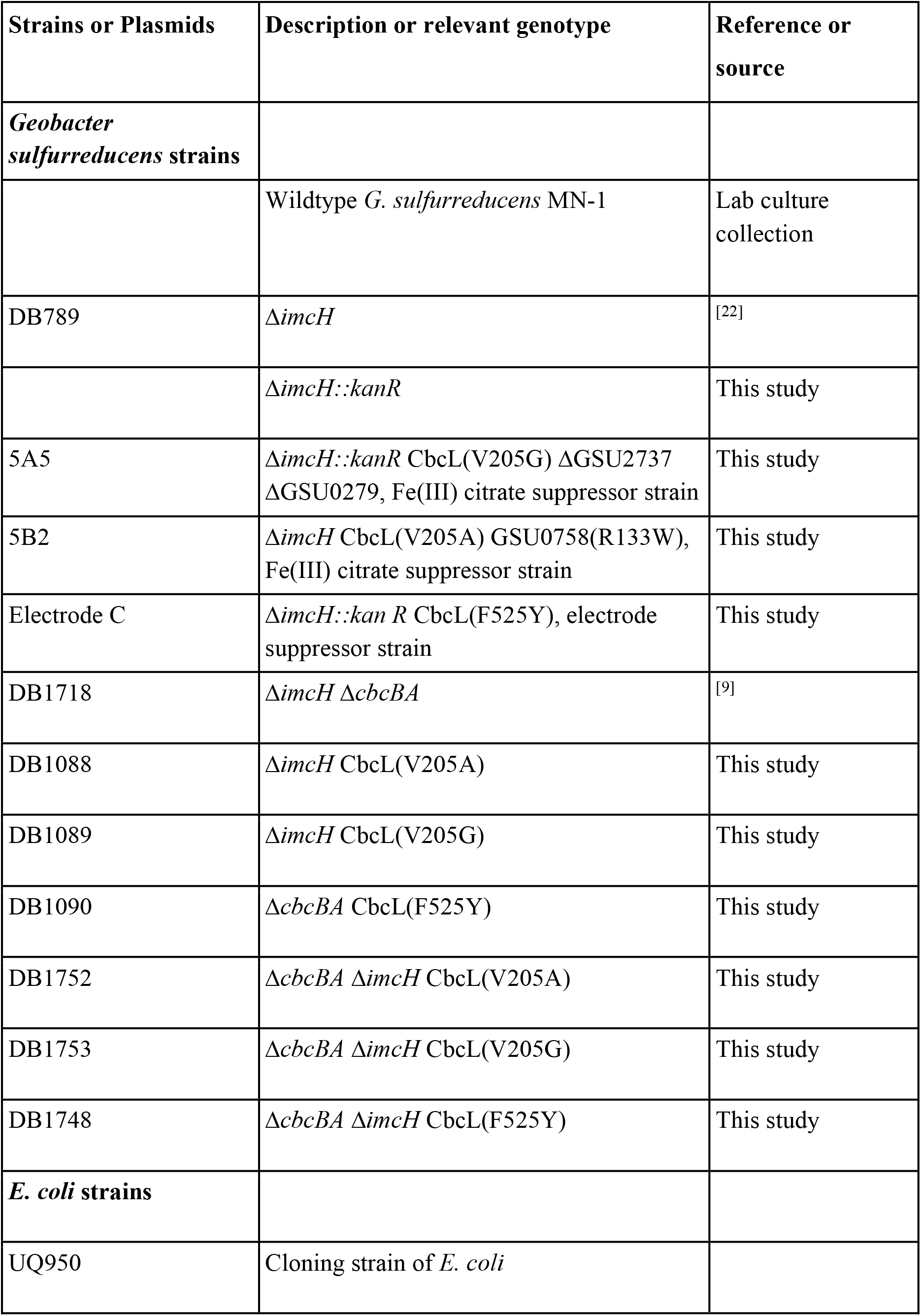
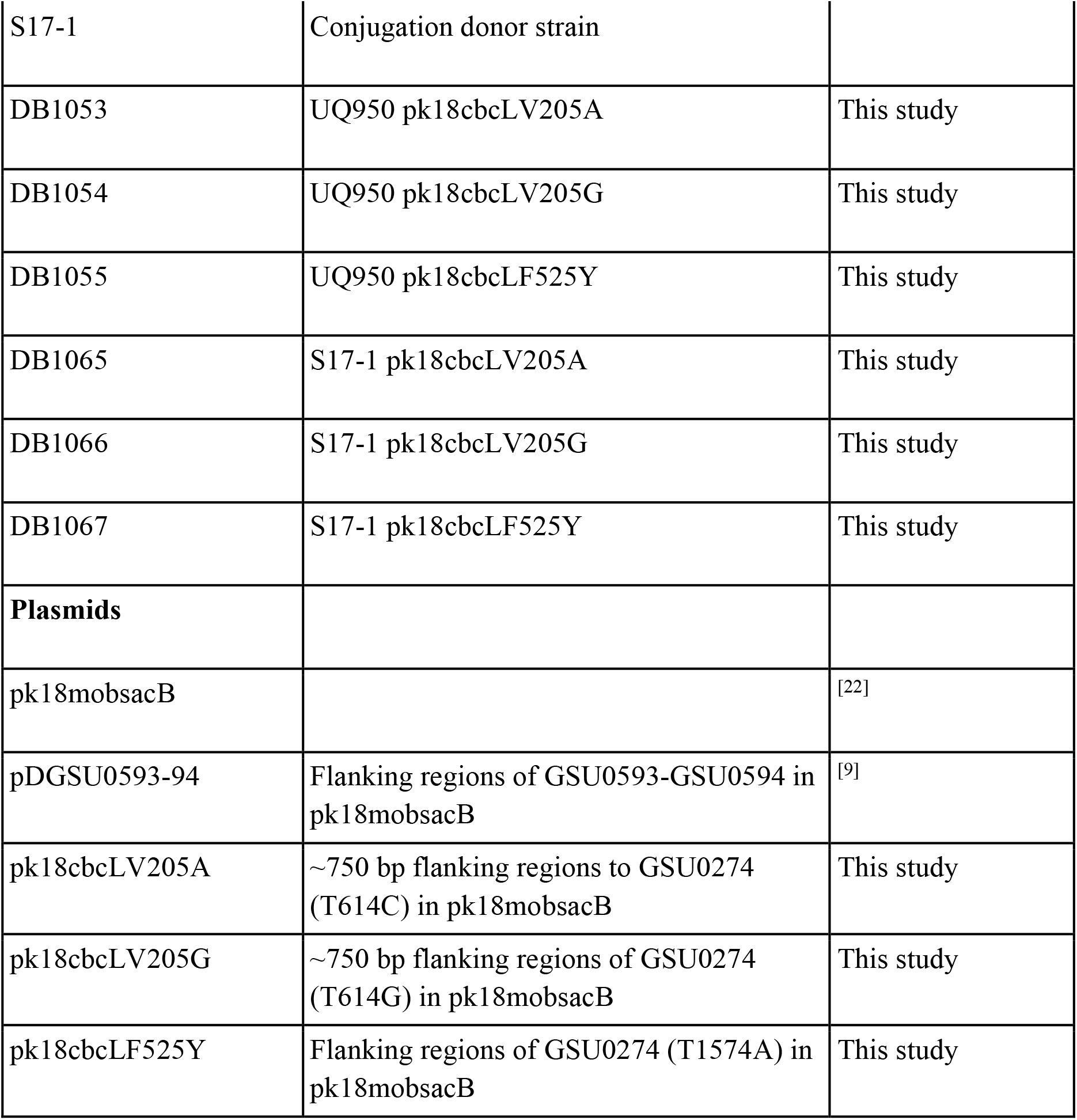
List of strains and plasmids used in this study

**Table 2:**
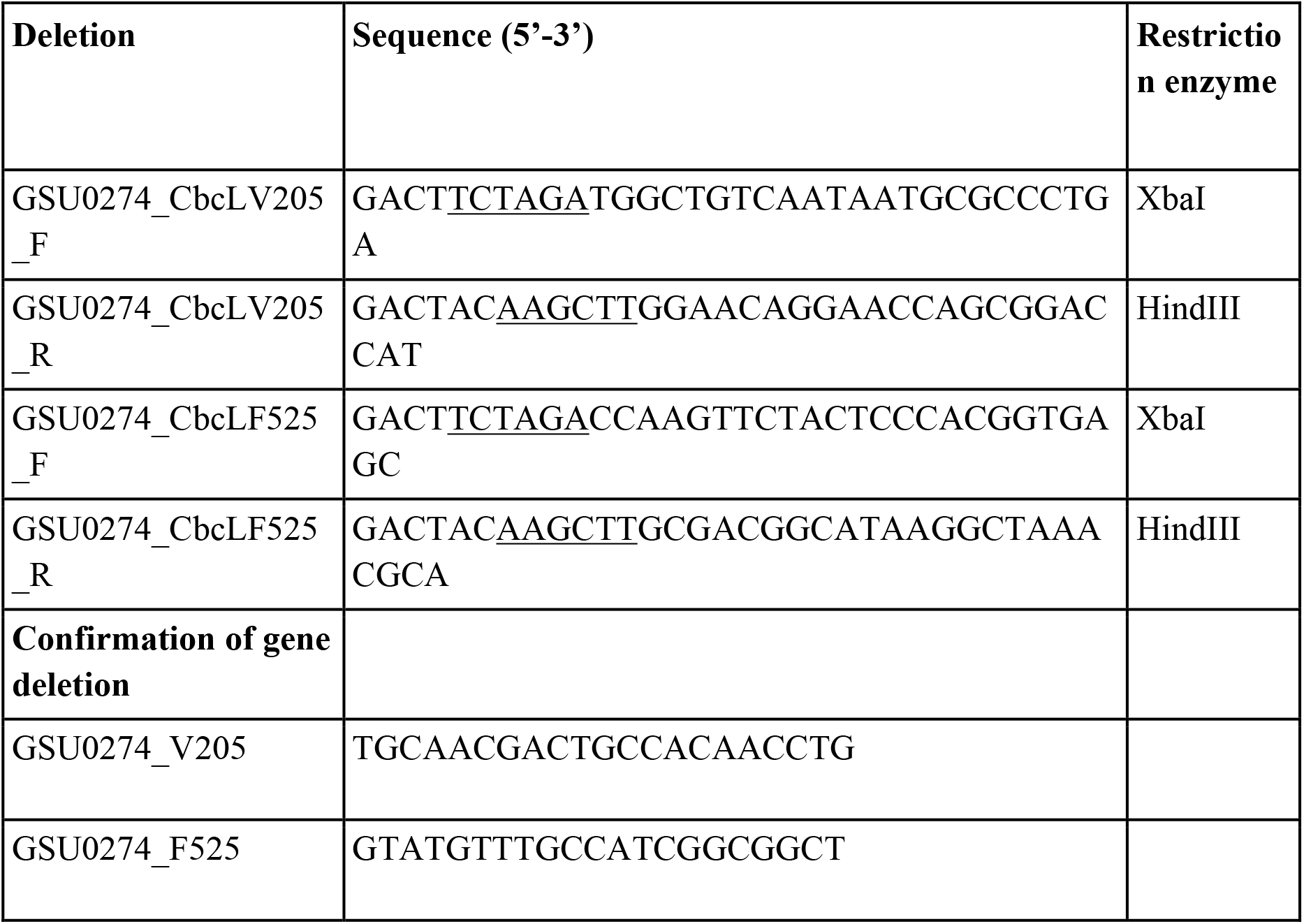
Primers used in this study.

